# Rare species perform worse than common species under changed climate

**DOI:** 10.1101/805416

**Authors:** Hugo Vincent, Christophe N. Bornand, Anne Kempel, Markus Fischer

## Abstract

Predicting how species, particularly rare and endangered ones, will react to climate change is a major current challenge in ecology. Rare species are expected to have a narrower niche width than common species. However, we know little whether they are also less able to cope with new climatic conditions. To simulate climate change, we transplanted 35 plant species varying in rarity to five botanical gardens in Switzerland, differing in altitude. For each species we calculated the difference in climate between their natural habitats and the novel climate of the respective botanical garden. We found that rare species had generally lower survival and biomass production than common species. Moreover, rare plant species survived less when the amount of precipitation differed more from the one in their natural range, indicating a higher susceptibility to climate change. Common species, in contrast, survived equally well under all climates and even increased their biomass under wetter or drier conditions. Our study shows that rarer species are less able to cope with changes in climate compared to more widespread ones, which might even benefit from these changes. This indicates that already rare and endangered plant species might suffer strongly from future climate change.

## INTRODUCTION

Understanding how species respond to a changing climate is one of the most important current challenges for ecologists (Chevin et al. 2010, Chessman 2013, Pacifici et al. 2015). Rare, already endangered species might be particularly vulnerable to climate change (Schwartz et al. 2006), and information in how they respond to changes in climate is crucial to target conservation and management efforts. For plants, the predicted changes in temperature and precipitation can have profound implications for their growth and survival. An increase of 1 to 2°C in the global mean surface temperature (IPCC 2014) along with a reduction in the average amount of precipitation, and the occurrence of more extreme events such as droughts, directly impact plants and change abiotic and biotic parameters. To survive climate change, plant populations may migrate to keep track of favorable environmental conditions, or they can also tolerate the new climatic conditions and adapt (Franks et al. 2014). Accordingly, many models predict that species will shift their ranges in response to climatic modifications (e.g. Bakkenes et al. 2002, Thomas et al. 2004). However, migration may be limited, e.g. by topographic boundaries such as mountains, the increasing fragmentation of our landscapes (Jump and Peñuelas 2005), or for species with a long generation time and low dispersal abilities (Aitken et al. 2008), and hence models hypothesize that a higher number of plant species will be threatened in a close future by the loss of climatically suitable areas (Thuiller et al. 2005). Therefore, tolerance to climate change might be of particular importance for plant populations.

One of the main hypothesis aiming to explain why some plant species are rare while others range widely is the niche-breadth hypothesis, which suggests that rare species are rare because they have a smaller niche breadth, i.e. they are less able to maintain viable populations across a range of environments, than more common species with a greater range size (Brown 1984, Slatyer et al. 2013). This hypothesis has achieved consistent support when quantifying the niche breadth based on the current distribution of species, suggesting that a positive relationship between range size and climatic niche breadth is a general pattern (Kambach et al. 2018). A species can have a large climatic niche because it consist of many locally adapted populations that each are adapted to different climatic conditions (Ackerly 2003) or because it consist of general-purpose genotypes that thrive in a wide range of environmental conditions through phenotypic plasticity (Baker 1965). Only the latter would enable plant populations to tolerate new climates when migration is hindered. However, we lack empirical knowledge on whether individuals of more common species are more tolerant to climatic variation, i.e. whether they have a larger fundamental niche due to general-purpose genotypes, than more rare and endangered species do. This information is crucial if we want to forecast the future composition of plant communities and to detect species that are particularly sensitive to climate change. Answering this question requires experimental approaches with many plant species (van Kleunen et al. 2014), however, empirical assessments of the fundamental climatic niches are scarce.

In this study, we tested the response of 35 plant species differing in rarity from rare and endangered to widespread species, to different climatic conditions. We used an altitudinal gradient in Switzerland, with a dryer and warmer climate at low altitudes and a wetter and colder climate at higher altitudes, to simulate climate change (Körner 2007). By transplanting the 35 plant species to five different botanical gardens along an altitudinal gradient, we were able to follow their survival and performance under various climatic conditions, which differed from the climatic conditions of their natural range. Using this experimental multi-species multi-site approach, we addressed the following specific questions: (i) Across different climatic conditions, do rare and common plant species generally differ in their survival and performance? (ii) Do rare and common plant species respond differently to changes in climatic conditions? We hypothesize that all species should perform best when the climatic conditions match the ones of their natural range. However, given that species with a small range size might also have a narrower fundamental niche width than more widespread species (Brown 1984), we expect individuals of rare species to be less tolerant to changes in climatic conditions, putting them at an even higher risk of extinction with climate warming.

## METHODS

### Plant species and experimental design

We used 35 plant species from 16 plant families (see Table S1 in Supporting Information). Twenty-four of those were rare species with a conservation priority in Switzerland (Moser et al. 2002), and 11 of them were common species which are widespread in Switzerland. Seeds of rare plant species were collected in the wild (one population per species) in Switzerland. Seeds of common species were collected in the wild or obtained from commercial seed suppliers (Rieger-Hofmann GmbH, Germany and UFA Samen, Switzerland). In March 2012, we germinated the seeds and planted 50 seedlings per species individually into 2-L pots filled with potting soil. Plants were then placed in a common garden (Muri near Bern, Switzerland) where they grew for another two months. In May 2012, we measured plant height to account for initial size differences. In June 2012 we transported the plants to five Botanical Gardens differing in altitude and climatic conditions (Table 1). In each garden, we placed 10 pots per species (occasionally less, Table S2) and distributed them randomly into garden beds. In early summer 2013 we recorded the survival of the plants and collected aboveground biomass. Since watering happened only in case of severe drought, we can assume that the observed differences in plant growth between the gardens is due to differences in precipitation and temperature and is not biased by the care taken by the botanical gardens.

**Table 1.** Location, altitude and climatic conditions of the five botanical gardens.

| Botanical garden | Coordinates<br>(CH1903) | Altitude (m) | Average annual<br>precipitations (mm) | Average annual<br>temperature (°C) |
| --- | --- | --- | --- | --- |
| Basel | 610797 - 267566 | 269.4 | 787.3 | 9.48 |
| Geneva | 500516 - 120219 | 372.2 | 909.5 | 9.53 |
| Pont-de-Nant | 500516 - 120219 | 1262.9 | 1451.1 | 5.98 |
| Champex | 574742 - 97996 | 1532.6 | 1376.9 | 4.19 |
| Schynige Platte | 636229 - 166947 | 1963.7 | 1630.6 | 1.61 |

### Rarity and climatic variables

To obtain a continuous measure of plant rarity we used the range size of each species in Switzerland. Range size was expressed as the number of 10 × 10 km grid cells occupied by a given species in Switzerland (data provided by Info Flora). We used range size in Switzerland because a continuous measure of European range sizes for our species is not available yet. Nevertheless, for a subset of 21 species for which European range size is available, Swiss and European range size were positively correlated (r = 0.508, p < 0.001, Text S1).

For each species we calculated climatic values, which characterize the climatic conditions in the natural range of a species in Switzerland. We calculated the mean annual temperature and mean annual level of precipitation per species (Table S1) by extracting climatic information at all known locations of the species in Switzerland using precise coordinates (for complete details on the climate data, see (Zimmermann and Kienast 1999). For each botanical garden, we also extracted the mean annual temperature and mean annual level of precipitation (Table 1).

To define the difference in climate between a botanical garden and a species’ natural range, we calculated the temperature and precipitation differences by subtracting the climatic value of a species range from the climatic value of a botanical garden. A negative value of a precipitation or temperature difference indicates that the climate is dryer or colder, respectively, in a botanical garden than in the species natural range. The range size of our species was not related to the mean altitude (r = 0.01, p = 0.95) and the mean temperature (r = −0.08, p = 0.64) of their natural range. Range size was positively related to the mean annual level of precipitation (r = 0.40, p = 0.02).

### Statistical analysis

To test whether species with a larger range size also occurred in a wider range of climates (i.e. whether they also have a larger climatic niche) we correlated range size with the difference between the maximum and the minimum value of temperature and precipitation of the species natural ranges. To test whether rare and common species generally differ in their survival and aboveground biomass production, we used generalized linear mixed effects models (*glmer)* with a binomial error distribution and linear mixed effects models (*lmer*) using the lme4 package (Bates *et al*. 2014) in R (R Core Team 2014), with the range size of the species as fixed term, the species identity nested into plant family (to account for taxonomy), and the botanical garden where the plants grew, as random factors. We also included the initial height of the plants as covariate, to control for initial size differences.

To test whether rare and common species respond differently in terms of their survival and aboveground biomass production to climatic differences, we used range size, temperature difference, precipitation difference, and the interaction between range size and climatic differences as explanatory variables. We also included the quadratic terms for the climatic differences as we expected a hump-shaped relationship with an optimum at a climatic difference of 0 (i.e. where the climatic conditions in a garden match the ones of a species natural range). Further, we included the interaction between the quadratic terms for the climatic differences and the range size of the species. Although the climatic variables ‘temperature difference’ and ‘precipitation difference’ were correlated with each other (r = −0.64, p < 0.001), both explained a significant part of the variation and were both kept in the model. We simplified the full models by removing non-significant terms and we determined significances using likelihood-ratio tests comparing models with and without the factor of interest. Non-significant linear terms were kept when the corresponding interaction and quadratic terms were significant. We log-transformed the biomass data and scaled all continuous variables to means of zeros and standard deviations of one for an easier interpretation of the model estimates.

## RESULTS

Range size strongly correlated with the species temperature and precipitation niche width, i.e. with the difference between the maximum and the minimum temperature (r = 0.83, p < 0.001), respectively precipitation (r = 0.78, p < 0.001) in the natural range. This confirms that more widespread species occur in a wider range of climatic conditions than rarer species. Overall, species with a larger range size survived better (Chi^2^ = 3.88, p = 0.049) and produced more aboveground biomass (Chi^2^ = 17.5, p < 0.001, Fig. 1) than rarer species.

**Figure 1.**
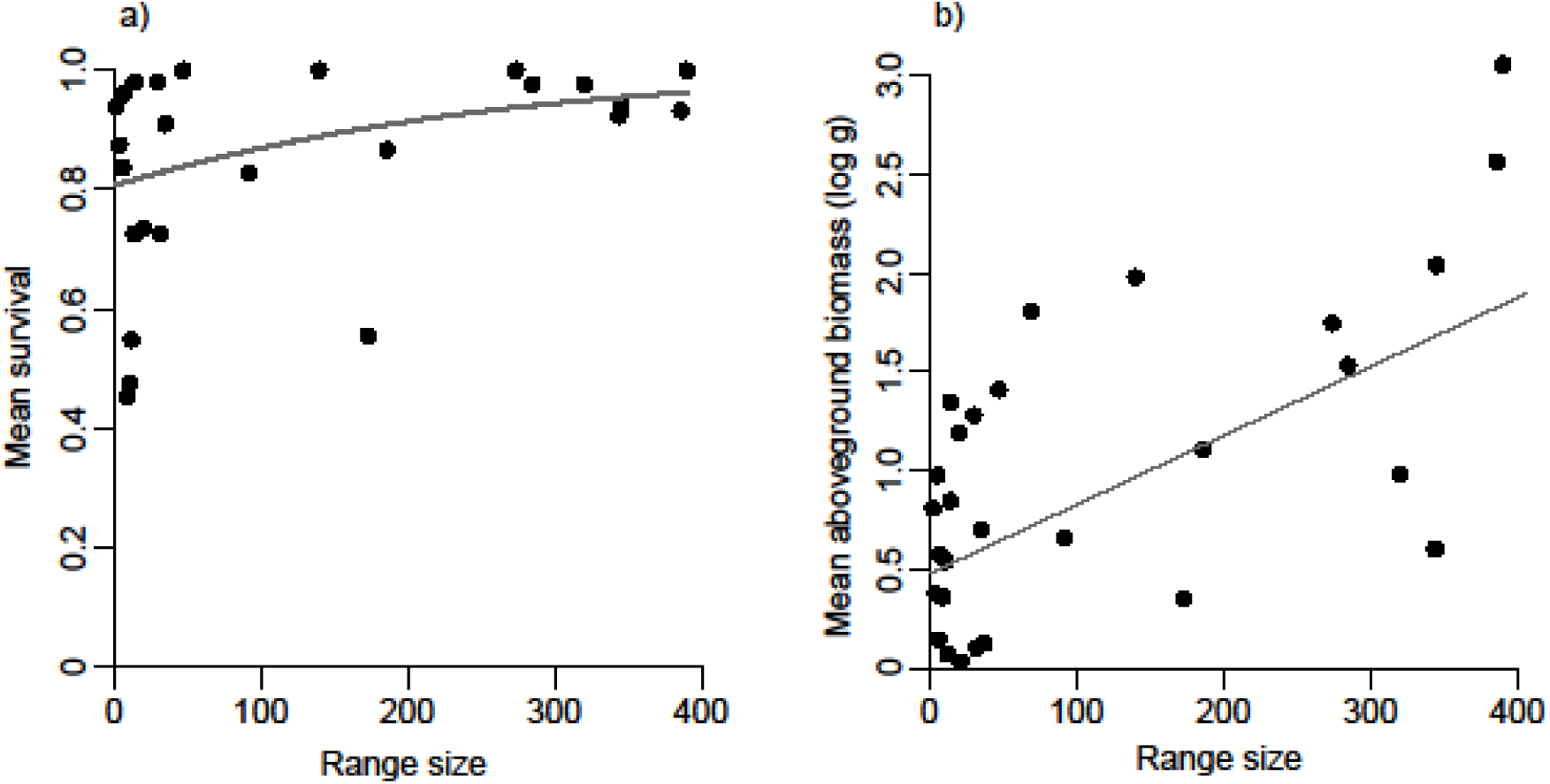
Effect of range size on a) mean survival, and b) mean aboveground biomass (expressed in g on a log-scale) for 35 species planted to five botanical gardens. Each point represents the mean biomass or survival per species, the line is obtained from the predicted values of the models. Range size is calculated as the number of 10×10km grid cell occupied by a given species in Switzerland. The curved line describing the relationship between range size and survival is obtained from the transformation of the binomial survival data into a continuous distribution of the probability of survival.

Survival was highest at low precipitation differences, i.e. when the climatic conditions of a garden were most similar to the ones of a species natural range. This effect was only driven by rare plant species, whose survival decreased when the amount of precipitation in a garden differed from the one of their natural range. In contrast, more common species were hardly affected by differences in precipitation, maintaining a high average survival in all botanical gardens (significant range size x squared precipitation difference interaction, Table 2, Fig. 2a).

**Table 2.** Results of a linear mixed effects model and a generalized linear mixed effects model testing for an effect of range size, temperature difference between natural sites and botanical garden (Δ Temperature), precipitation difference (Δ Precipitation), the quadratic terms of Δ Temperature and Δ Precipitation, and their interactions on biomass production and plant survival of plants of 35 species planted to five botanical gardens. We removed all non-significant terms, unless the respective quadratic or interaction term was significant. All explanatory variables are scaled. The parameters of the main factors that were present in significant interactions were derived from models where all higher order interactions were removed.

| *Fixed terms* | *Biomass* |  |  | *Survival* |  |  |
| --- | --- | --- | --- | --- | --- | --- |
|  | estimate | Chi<sup>2</sup> | p-value | estimate | Chi<sup>2</sup> | p-value |
| Range | 0.24 | 11.2 | <0.001*** | 0.14 | 1.91 | 0.17 |
| $\Delta$ Temperature | -0.05 | 1.59 | 0.206 | -0.01 | 2.95 | 0.086 |
| $\Delta$ Precipitations | -0.23 | 11.6 | 0.03* | -0.16 | 0.05 | 0.831 |
| $\Delta$ Temperature<sup>2</sup> | -0.13 | 67.8 | <0.001*** | -0.55 | 32.2 | <0.001*** |
| $\Delta$ Precipitations<sup>2</sup> | 0.04 | 0.58 | 0.446 | -0.27 | 8.85 | 0.003** |
| Range x $\Delta$ Temperature | - | - | - | - | - | - |
| Range x $\Delta$ Precipitations | 0.01 | 15 | <0.001*** | - | - | - |
| Range x $\Delta$ Temperature<sup>2</sup> | - | - | - | - | - | - |
| Range x $\Delta$ Precipitations<sup>2</sup> | 0.09 | 26.2 | <0.001*** | 0.2 | 4.54 | 0.033* |
| *Random terms* | *Variance* |  |  | *Variance* |  |  |
| Species | 0.265 |  |  | 1.88 |  |  |
| Family | <0.001 |  |  | <0.001 |  |  |
| Botanical Garden | 0.016 |  |  | 0.924 |  |  |

**Figure 2.**
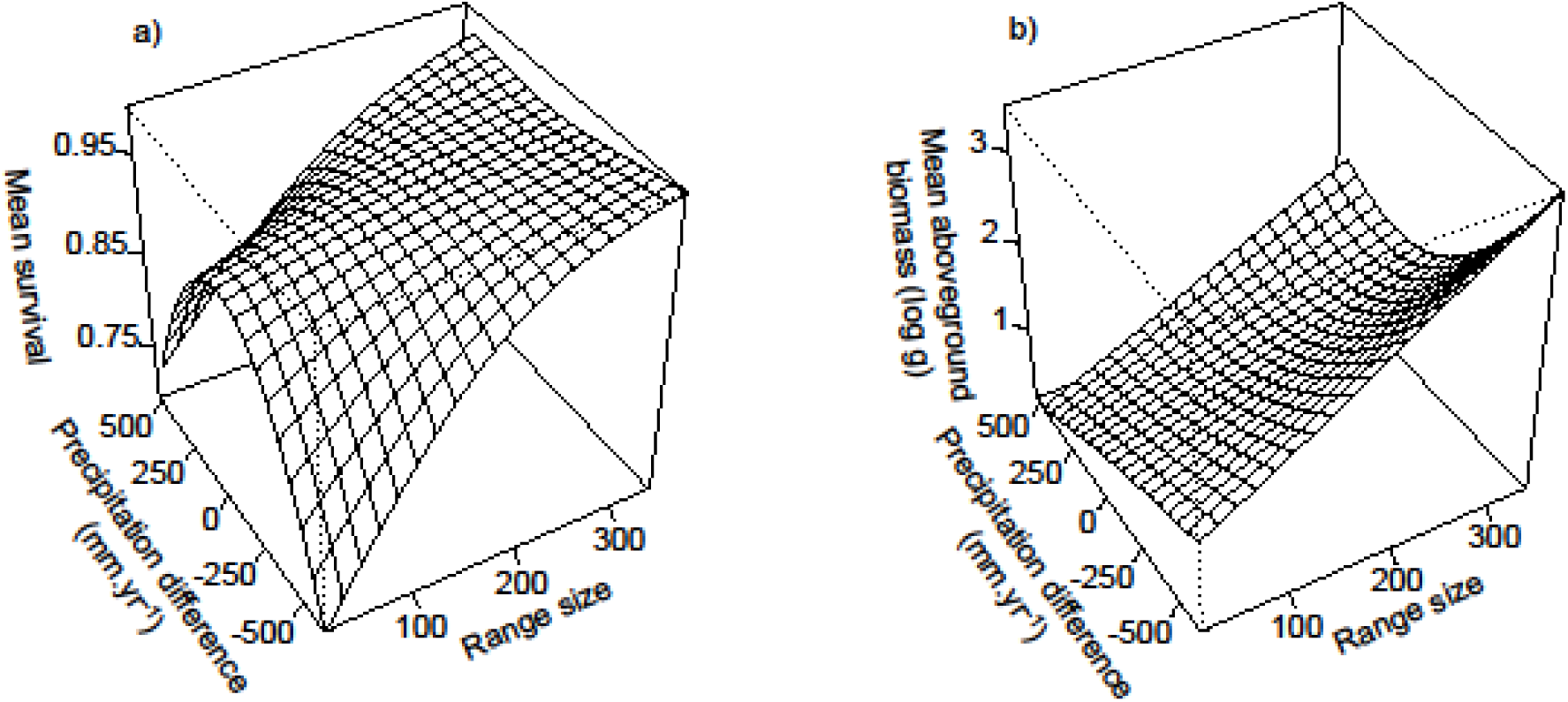
a) Survival and b) biomass production of 35 species in relation to precipitation difference between natural range size and botanical garden. The surfaces represent the predicted survival, respectively biomass, from the model. Biomass is expressed in g on a log-scale. A negative precipitation difference (mm year^-1^) indicates that the conditions in a garden are dryer than the ones in a species natural range.

Aboveground biomass of rarer species was hardly affected by differences in precipitation between a botanical garden and the species natural range. Common species, however, produced more biomass when the conditions were drier - and thus sunnier - and when the conditions were wetter than in their natural range (Table 2, Fig 2b). This indicates that more common plant species are able to plastically increase their biomass in these conditions whereas rarer plant species are less plastic and show a relatively stable biomass production.

Overall, survival and biomass production was lowest when the temperature in a botanical garden deviated most from the mean temperature of a species natural range (significant squared temperature difference effect, Table 2, Fig. 3), and this did not differ for rare and common species.

**Figure 3.**
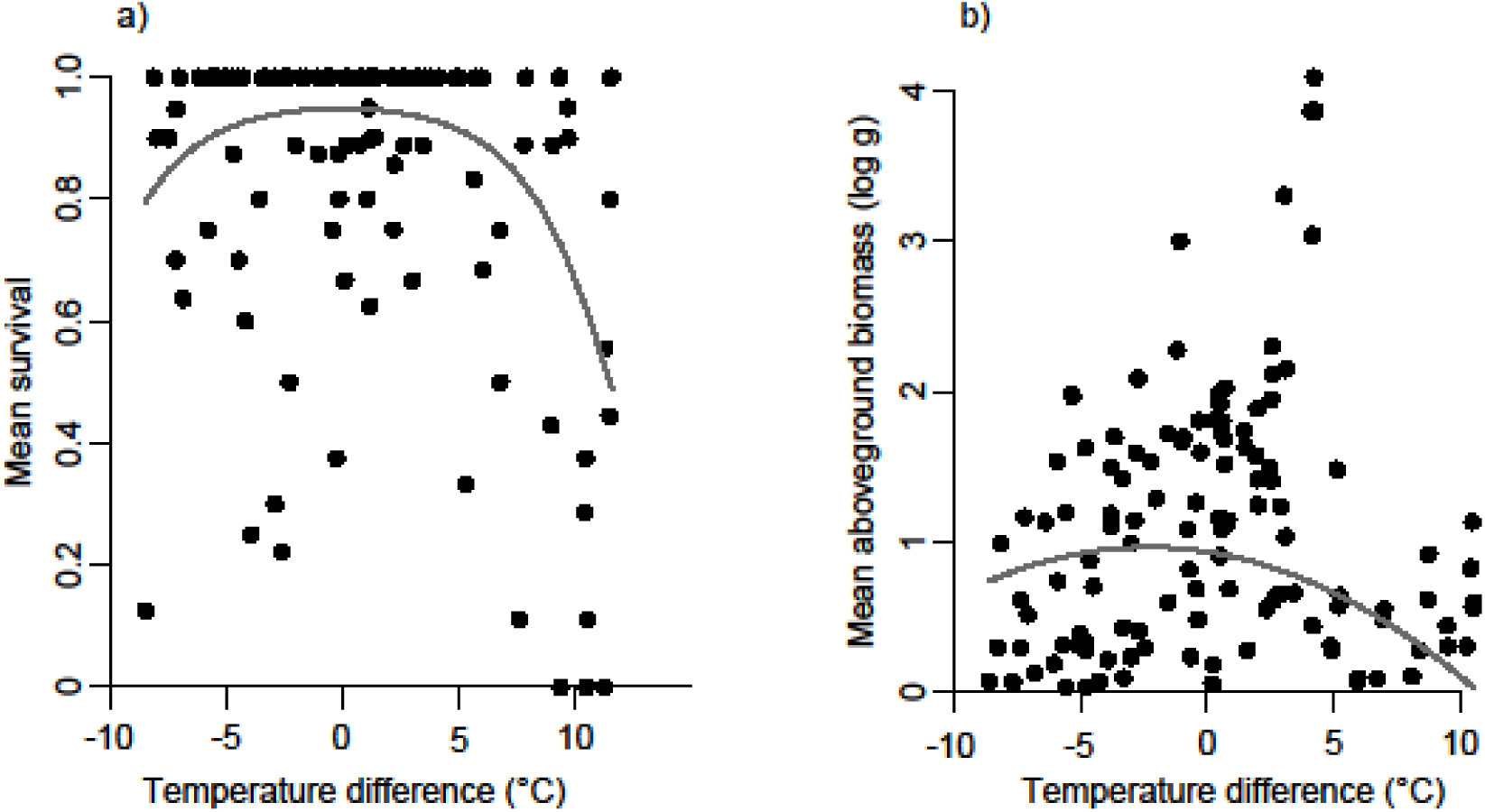
Effect of the temperature differences (°C) on a) mean survival and b) mean aboveground biomass of 35 species planted in five botanical gardens. Each point represents the average aboveground biomass (in g on a log-scale) or survival per species per garden in 2013. The line is obtained from the predicted values of the models. To represent the effect of temperature difference, we fixed the value of precipitation difference to its mean when calculating the predicted values of the models.

## DISCUSSION

### Rare plant species are less tolerant to changes in climate than common plant species

Among the most important hypotheses explaining species rarity and commonness is the niche breadth hypothesis, which predicts that species that are able to maintain populations across a larger set of environmental conditions can achieve larger geographic ranges than species with narrow ecological niches (Brown 1984). Studies relating the range size of species to their realized niches supported the predicted pattern (Kambach et al. 2018). However, whether this means that more common species also consists of individuals, which generally have larger fundamental niches (general-purpose genotypes) than have rare species, and therefore which have a higher ability to cope with changing climatic conditions, remains unknown. In our experiment, plant species generally survived better and had a greater biomass when the mean annual temperature of the botanical gardens was similar to the one they experience in their natural range (Fig. 3), reflecting the existence of a climatic niche due to physiological limitations, which is a key assumption for predicting the impact of climate change on species distributions (Pearman et al. 2008, Petitpierre et al. 2012). Similarly, plants survived better when the mean annual precipitation mirrored the one from their natural range, however, this was only driven by rare plant species, which suffered from differences in precipitation (when conditions where either dryer or wetter than the ones at their origin). In contrast, more common species were not affected by precipitation differences, and showed a similarly high survival at all precipitation levels, independent of the ones of their origin (Table 2, Fig. 2a). Our results demonstrate that rarer species do indeed have a smaller fundamental niche in terms of precipitation, i.e. a lower climatic tolerance due to physiological limitations, than more common species. Since climate change is expected to increase wet and dry extreme events (Knapp et al. 2008) this suggests that species, which are already threatened under the current climate will suffer most from the effects of climate change.

Widespread species are likely to experience a larger range of ecological and climatic conditions within their range (Gaston 2003). Indeed, a larger niche width – based on the current distribution of a species – seems to be a general pattern in widespread species (Slatyer et al. 2013, Kambach et al. 2018), and was also supported by our data (positive correlation between range size and the climatic width). A species can accrue a larger niche breadth because it consists of many locally adapted populations (Olsson et al. 2009) which partition the broad climatic tolerance exhibited by the species as a whole. Moreover, species can be composed of phenotypically plastic genotypes, general-purpose genotypes or individual generalists that perform well under a large range of environmental conditions (Baker 1965, Ackerly 2003). Although in our experiment we cannot entirely disentangle the factors leading to a higher climatic tolerance in common species, the fact that we found this pattern by placing only a few individuals into the different botanical gardens indicates that widespread species are more likely to be comprised of individual ‘generalists’. However, to fully understand the influence of broad tolerance and microevolution on niche width, experiments simultaneously comparing climatic tolerance of many species, populations per species, and genotypes per population are needed.

In contrast to results on survival, aboveground biomass production of rarer species hardly changed in response to differences in precipitation. More common species, however, increased their biomass particularly when the amount of precipitation was lower than in their natural range (Table 2, Fig. 2b). Possibly, a dryer climate implies a higher number of sunny days and therefore more favorable conditions for plant growth. More common species therefore seem to be more able to plastically increase their biomass under favorable growing conditions, whereas rarer species seem to be less able to change their phenotypes in response to environmental variation. When precipitation was higher than in their natural range, more common species were also able to increase their biomass. This plastic response in more widespread species indicates that, in addition to maintaining generally high survival under different climatic conditions, widespread species were able to take advantage of both drier and wetter conditions. Widespread species have also been shown to be better able to take advantage of an increase in nutrient availability than rare species (Dawson et al. 2012) and, compared with species confined to river corridors, to better take advantage of benign conditions of non-river corridor conditions (Fischer et al. 2010). Our study therefore adds additional evidence that widespread species might be widespread as they are able to take advantage of favorable climatic and environmental conditions than species of small range size, and that this is a general pattern. Under future climate change, with a predicted increase in extreme precipitation events (Easterling et al. 2000), our results indicate that more common species might better take advantage of the changing climatic conditions and potentially outcompete rarer species. This calls for developing measures to support rare species.

In most cases, widespread species experience a wider range of climatic conditions in their natural ranges than species with a more restricted range size. Therefore, the mean altitude, mean annual precipitation and mean temperature of the 11 species common in Switzerland was intermediate among those of the 25 rare species, some of which only occur in alpine or lowland regions (Figure S1). This reduced the range of data points in climatic differences for common species and might have affected extrapolations of our models at the extreme ends of climatic differences. To control for such potential bias, we analyzed a subset of our data by keeping only those rare species that occur within the same climatic range than our common species (Table S3). This analysis confirmed the effects of climatic differences and their interaction with range size found for the whole dataset, which suggests that our finding that more widespread species have a wider climatic tolerance than rarer ones is robust.

Experimental tests of environmental tolerance of multiple plant species as the one we present here, and particularly of rare and common native species, are extremely rare (Slatyer et al. 2013). A few studies assessed the tolerance to different germination conditions (fundamental germination niche widths) of rare and common plant species and found either a positive (Brändle *et al*. 2003; Luna *et al*. 2012), negative (Luna and Moreno 2010) or no relationship with range size (Thompson and Ceriani 2003, Gaston and Blackburn 2007). Our results therefore highlight that plant rarity is related to the fundamental climatic niche of species, and calls for a more differentiated view when predicting the future distribution of different species to climate change.

### Rare plant species have lower survival and lower biomass than common plant species

Why some species are rare while others are common has fascinated ecologists for decades (Brown et al. 1996, Webb and Gaston 2003). Differences in species characteristics have repeatedly been suggested to explain the distribution and abundance of plant species in nature (Murray et al. 2002, Kempel et al. 2018). In our study, overall, rare plant species showed lower survival and lower biomass production than common plant species. This variation in the intrinsic general performance of plants could be a major driver of rarity and commonness at large spatial scales. Lower biomass of rare species has also been found in other studies (Murray et al. 2002, Lavergne et al. 2003, Cornwell and Ackerly 2010, Dawson et al. 2012, Kempel et al. 2018) and indicates that rare species have slower growth rates (Cornelissen et al. 2003), a trait that is often attributed to slower nutrient uptake and hence lower competitive ability in productive habitats (Grime 1977). By using a continuous gradient of rarity and commonness with many species originating from different habitats, our approach suggests that a positive relationship between plant performance and plant range size is a general pattern. Future studies that take various aspects of rarity into account, including small and large populations of plant species differing in range size, are needed to ultimately test whether a lower general performance of species of small distribution range is a result of small population sizes and hence reduced genetic diversity (Leimu et al. 2006), or whether generally lower general fitness of such species is responsible for their small distributional ranges.

### Conclusion

Using a large number of plant species differing in their range size in Switzerland, we provide experimental evidence that more widespread species indeed have larger climatic niches than rarer species. We showed that rare species not only have generally lower survival and biomass production than more common species but that they are also more susceptible to a changing climate. On the contrary, more widespread species survived equally well under all climates and could even take advantage of favorable growing conditions by plastically increasing their biomass. Our multi-species experiment suggests that this is a general pattern. We conclude that already rare and endangered plant species have a lower climatic tolerance than more widespread species and might suffer strongly from the forecasted climatic changes.

## ACKNOWLEDGEMENTS

We thank Adrian Möhl and Christine Föhr for collecting seeds in the natural populations; the teams of the Botanical Gardens of Basel, Champex, Geneva, Pont-de-Nant and Schynige Platte, for hosting the experiment and for their support; the numerous field assistants who helped harvesting the plants; Niklaus E. Zimmermann for providing the original climatic data; Eric Allan and Santiago Soliveres for their comments on an earlier draft of this manuscript. This study was supported by the Federal Office of the Environment, Switzerland.

## FIGURES LEGENDS

**Table S1.**
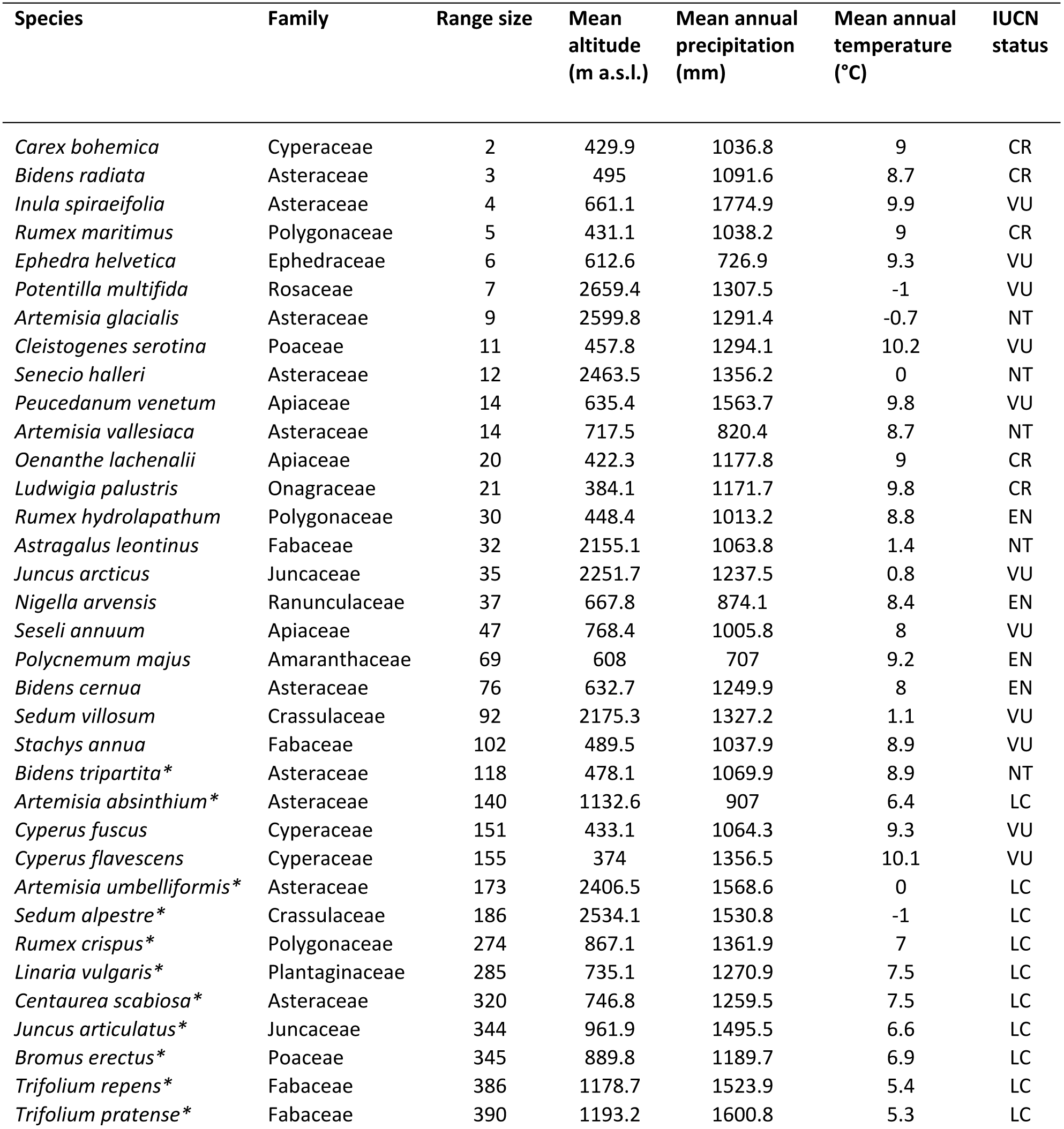
List of the 24 rare and 11 common species (indicated by *) studied in this experiment, including their plant family, range size in Switzerland (number of 10×10 kilometers grid cells occupied by a species in Switzerland, see Methods), mean altitude, mean annual amount of precipitation and temperature of the species natural range, and the IUCN category of threat in Switzerland (LC: Least Concern; NT: Near Threatened; VU: Vulnerable; EN: Endangered; CR: Critically Endangered).

**Table S2.**
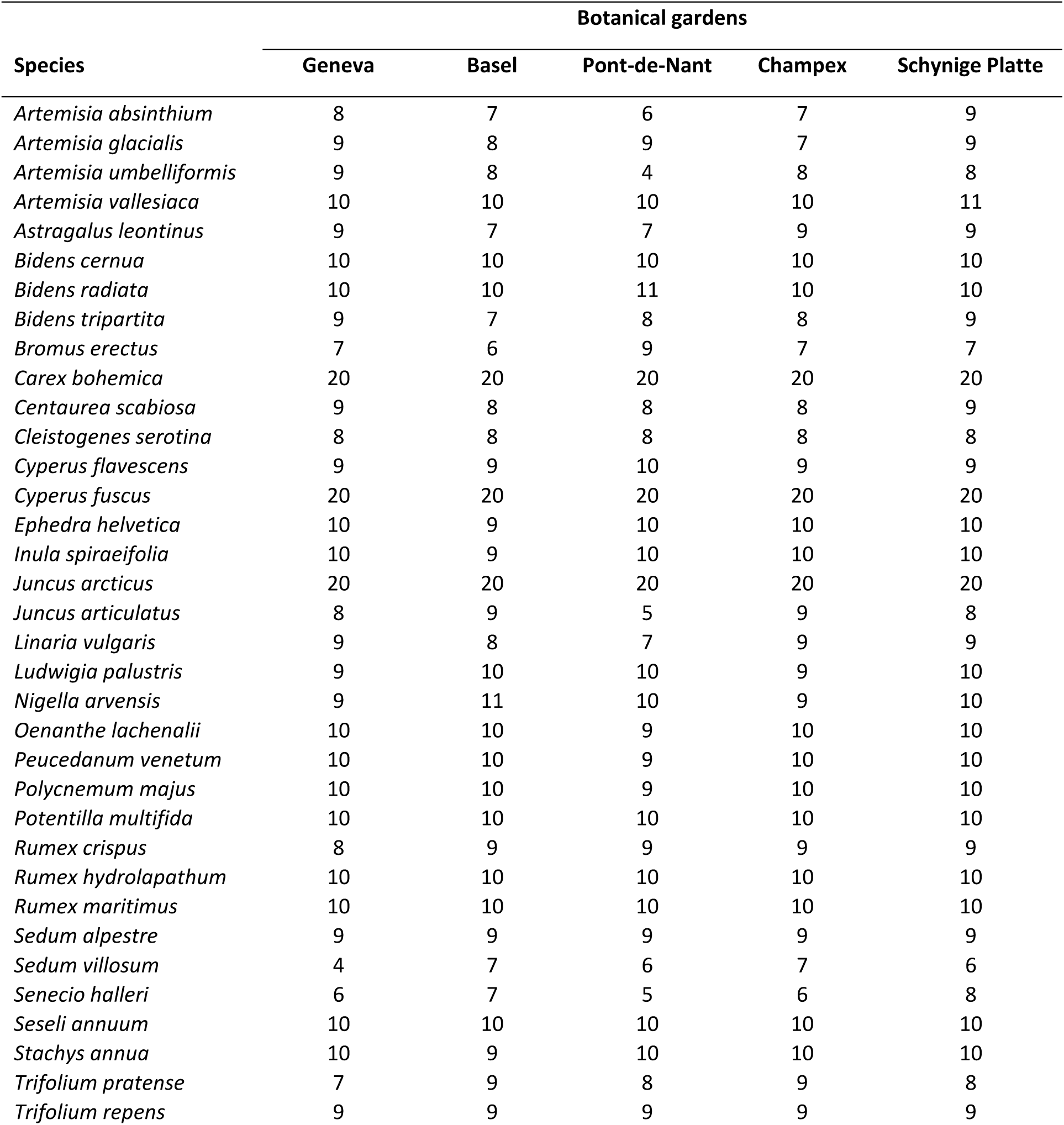
Number of plants per species grown in each botanical garden.

**Table S3.** Effect of climatic differences on the biomass production and the survival of a subset of 31 species. The rare species used in this experiment naturally occur in a wider range of climatic conditions than the common species used in this experiment (Fig. S4). We re-analyzed our data with a dataset including all the common species and a subset of 20 rare species, keeping only those which occur inside a precipitation range of 900 to 1600 mm.yr^-1^. We considered the precipitation values to define this climatic range because it was the climatic variable which interacted with range size. The results did not differ qualitatively from the analysis of the entire dataset.

| *Fixed terms* | *Biomass* |  |  | *Survival* |  |  |
| --- | --- | --- | --- | --- | --- | --- |
|  | estimate | Chi<sup>2</sup> | p-value | estimate | Chi<sup>2</sup> | p-value |
| Range | 0.19 | 7.37 | 0.006** | 0.63 | 0.75 | 0.391 |
| Δ Temperature | -0.09 | 0.71 | 0.4 | 1.17 | 1.86 | 0.173 |
| Δ Precipitations | -0.28 | 5.73 | 0.017* | - | - | - |
| Δ Temperature<sup>2</sup> | -0.15 | 82.8 | <0.001*** | -0.58 | 30.8 | <0.001*** |
| Δ Precipitations<sup>2</sup> | 0.01 | 1.75 | 0.186 | -0.18 | 0.08 | 0.778 |
| Range x Δ Temperature | - | - | - | - | - | - |
| Range x Δ Precipitations | 0.05 | 11.4 | <0.001*** | - | - | - |
| Range x Δ Temperature<sup>2</sup> | - | - | - | - | - | - |
| Range x Δ Precipitations<sup>2</sup> | 0.11 | 30.1 | <0.001*** | 0.2 | 5.22 | 0.022* |
| *Random terms* |  |  |  |  |  |  |
|  | *Variance* |  |  | *Variance* |  |  |
| Species | 0.284 |  |  | 17.63 |  |  |
| Family | <0.001 |  |  | <0.001 |  |  |
| Botanical Garden | 0.018 |  |  | 1.818 |  |  |

**Text S1**. To test whether range size in Switzerland is correlated with the European range size of our study species, we used map-derived area estimates from the Atlas Europeae (Meusel *et al*. 1978) for the 21 species for which these maps were available. We assessed the number of pixels of a species European distribution and cross-referenced these using islands, for which the exact surface values are known. Range size in Europe was correlated with range size in Switzerland (r = 0.508, p < 0.001).

Meusel, H., Jäger, E. J., Rauschert, S. & Weinert, E. (1978). *Vergleichende Chorologie der zentraleuropäischen Flora. Bd. 2, Text u. Karten*. Gustav Fischer Verlag, Jena.

**Figure S1.**
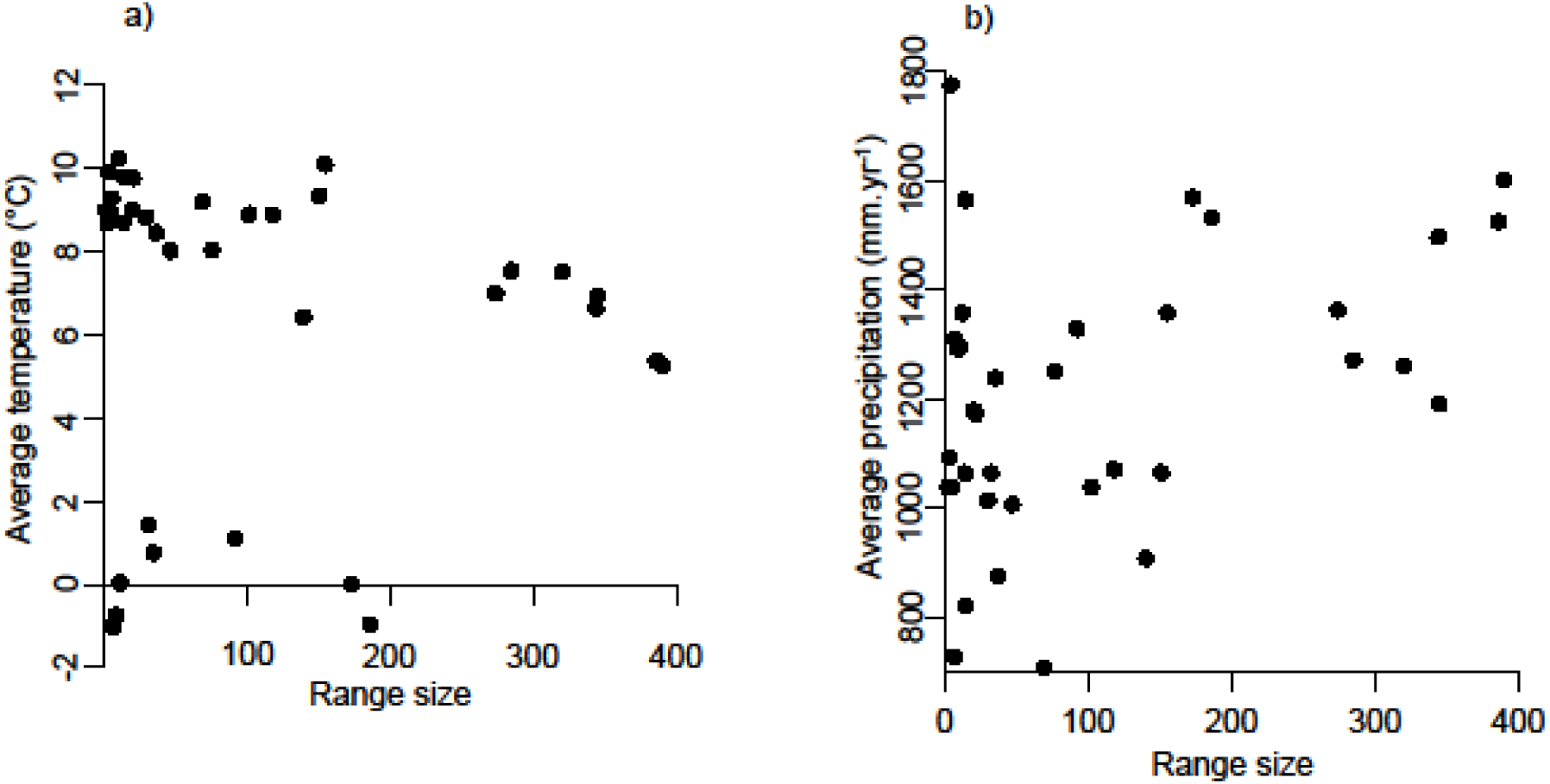
Correlations between a) mean temperature (°C) and b) mean annual level of precipitation (mm.year^-1^) in the natural range of our 35 species, and their range size. Common species showed more intermediate values than rarer species, although there was no correlation between range size and mean temperature (r = −0.08, p = 0.64), and the correlation between range size and mean annual precipitation (r = 0.40, p = 0.02) was not strong.

